# Evolutionary computing to assemble standing genetic diversity and achieve long-term genetic gain

**DOI:** 10.1101/2023.05.05.539510

**Authors:** Kira Villiers, Kai P. Voss-Fels, Eric Dinglasan, Bertus Jacobs, Lee Hickey, Ben J. Hayes

## Abstract

Loss of genetic diversity in elite crop breeding pools can severely limit long-term genetic gains, and limit ability to make gains in new traits, like heat tolerance, that are becoming important as the climate changes. Here we investigate and propose potential breeding program applications of optimal haplotype selection (OHS), a selection method which retains useful diversity in the population. OHS selects sets of candidates containing, between them, haplotype segments with very high segment breeding values for the target trait. We compared the performance of OHS, the similar method optimal population value (OPV), truncation selection on genomic estimated breeding values (GEBVs), and optimal cross selection (OCS) in stochastic simulations of recurrent selection on founder wheat genotypes. After 100 generations of inter-crossing and selection, OCS and truncation selection had exhausted the genetic diversity, while considerable diversity remained in the OHS population. Gain under OHS in these simulations ultimately exceeded that from truncation selection or OCS. OHS achieved faster gains when the population size was small, with many progeny per cross. A promising hybrid strategy, involving a single cycle of OHS selection in the first generation followed by recurrent truncation selection, substantially improved long term gain compared with truncation selection, and performed similarly to OCS. The results of this study provide initial insights into where OHS could be incorporated into breeding programs.

**Core Ideas:**

- We investigate potential uses for a haplotype-stacking strategy, optimal haplotype selection (OHS)
- Several selection strategies were compared in stochastic simulations of recurrent selection in wheat
- OHS maintained more useful diversity than optimal cross selection or truncation-based genomic selection
- Rates of gain from OHS are competitive in small populations
- One generation of OHS in a truncation selection program can increase short and long term genetic gain

**Plain language summary:** Breeders use selection strategies based on genetic and phenotypic information to choose parents that will improve agriculturally-relevant traits (eg. grain yield) in their progeny. Generally, this involves estimating breeding values (scores) for each candidate parent. This study investigated an alternative ‘haplotype stacking’ approach called Optimal Haplotype Selection (OHS), which instead estimates breeding values for each unique genome segment in the population, then selects a group of parents who, between them, carry the haplotypes with the highest estimated breeding value at each chromosomal segment. In simulations, OHS gives improvements close to existing methods when populations are small, and outperforms them in the long term (100+ generations). Using just one generation of OHS boosts the performance of other methods in the short and long term. Breeders might consider adopting haplotype stacking in their programs, once techniques to do so are established.

## 1 Introduction

A breeding program can be seen as a targeted reduction in the diversity of a population. Undesirable alleles are culled, and beneficial alleles increase in frequency. However, a breeding program with no diversity can produce no genetic gains. A significant challenge for elite plant breeding programs is to maintain or increase rates of gain over long time horizons without exhausting their breeding program’s genetic diversity.

Crop breeding programs are increasingly turning to genomic selection (GS) to increase genetic gains. GS involves ranking selection candidates (or candidate parents, for parent selection), on genomic estimated breeding values (GEBVs), which are calculated using genome-wide marker effects estimated from a phenotyped reference population (Meuwissen et al. 2001). Genomic selection can provide very high rates of gain in some species, as it enables early selection of breeding individuals and reduction in cycle times (Voss-Fels et al. 2019). The introduction of GS has boosted genetic gain in both animal and plant breeding programs (García-Ruiz et al. 2016; Guinan et al. 2023; Scott et al. 2021; Wolc et al. 2015; Crossa et al. 2014; Cooper et al. 2014; Bernardo 2021; Hickey et al. 2017).

An emerging strategy to integrate GS in crop breeding programs is rapid recurrent selection on GEBVs, especially in the population improvement component of a two-part breeding program (Gaynor et al. 2017). While recurrent truncation selection like this leads to rapid gains, it can cause dramatic loss of diversity and often leads to rapid inbreeding in the breeding pool (Lin et al. 2016; Cowling et al. 2019). In dairy cattle, for example, the rate of inbreeding has doubled as a result of the introduction of genomic selection (Makanjuola et al. 2020).

Potentially, the loss of diversity caused by truncation selection could be mitigated by introgressing genetically distant germplasm into the breeding program (Gorjanc et al. 2016, Allier et al. 2020, Sanchez et al. 2023). However, truncation selection is not well suited to introgression scenarios. Diverse or landrace germplasm, which may be carrying novel beneficial alleles, often lacks important agronomic traits compared to elite varieties that have been developed through generations of targeted selection. Under truncation selection, these mediocre GEBVs of diverse germplasm would result the diverse germplasm being culled early from the population. The value of novel alleles within the diverse germplasm are ‘masked’ by the GEBV calculation at the whole-genome level (Gorjanc et al. 2016). Truncation selection does not take into account any diversity measures that could compensate for this preference for elites.

Optimal contribution selection (OCS) is a strategy for balancing genetic gain and genetic diversity in animal and plant breeding programs (Wray and Goddard 1994; Meuwissen 1997; Woolliams et al. 2015; Lin et al. 2017; Gorjanc et al. 2018). In OCS, breeders weight two objectives according to their priorities: the minimisation of genetic relatedness, and the maximisation of genetic gain. The technique advises the proportions of genetic contribution from each candidate parent to best achieve the weighted objectives, or, with slight adaptation, suggests mate allocations among the candidate parents (“optimal cross selection”, Kinghorn 2011). OCS was designed for animal breeding and is seeing increasing use in plant breeding. Different plant breeding program designs based on OCS have been proposed (Cowling et al. 2017; Cowling et al. 2019), and it has been applied in forestry (Kerr et al. 2015), common bean breeding (Saradadevi et al. 2021), and spring canola breeding (Cowling et al. 2023).

To take advantage of genetic marker data, OCS can be carried out using GEBVs as the measure of genetic value and the genomic relationship matrix (GRM) as the measure of relatedness. Several variations on OCS exist. Genomic Mating (Akdemir & Sánchez 2016) and Usefulness Criterion Parental Contribution (UCPC, Allier et al. 2019) incorporate the expected variance in progeny value into the computation, to slightly improve results. Genomic OCS (GOCS, Tiret et al. 2021) adds a term to the GRM to improve genetic gain when using non-inbred populations. Recurrent application of OCS itself can provide rates of gain in line with or outperforming recurrent truncation selection within 15-20 generations of simulation, after a slight lag for the first few generations (Akdemir & Sánchez 2016, Sanchez et al. 2023).

While GS and OCS use whole-genome ratings to guide selection, selection methods could also be designed around the concept of collecting beneficial alleles into one genotype. Stacking, “gene pyramiding” or “genotype building” approaches in their direct form have not been widely applied in plant breeding (Han et al. 2017). In stacking approaches, genetic gain relies on beneficial recombination events causing the accumulation of good alleles or good segments in the same line (Dekkers and Hospital 2002; Servin et al. 2004). The minimum population sizes to guarantee fixation of desirable alleles or segments grow exponentially with the number of alleles or segments being targeted (Hospital 2003; Servin et al. 2004).

Therefore, stacking approaches do not seem suited to use together with GS, which uses thousands of markers. However, bundling markers into chromosome segments and selecting on chromosome segment haplotypes, as demonstrated for animal breeding by Kemper et al. (2012), could make this kind of selection feasible. This approach assumes every chromosome segment contains at least one mutation affecting the trait. It is effectively a version of gene-pyramiding adapted to a quasiinfinitesimal model of quantitative trait variation (many mutations of small effect spread across the genome). The optimal haplotype stacking (OHS) approach described by Kemper et al. (2012) was to select a set of candidates whose pool of haplotype segments could be stacked into the best “ultimate genotype”, the highest-scoring genotype that can be constructed out of the segments in the population.

A number of authors have considered stacking approaches for plant breeding applications. The optimal haploid value (OHV) (Daetwyler et al. 2015) of a candidate is the GEBV of the best doubled haploid that could be produced from that candidate out of its diploid marker block haplotypes. Selection on the OHV of candidates in a doubled-haploid-based breeding program outperformed selection on candidate’s GEBV for both genetic gain and diversity retention (Daetwyler et al. 2015). Variants of OHV selection exist that extend the method by calculating a score for each potential mating pair or each potential selection of candidates.

Han et al. (2017) treated all alleles as either desirable or undesirable in order to define the predicted cross value (PCV) as the probability that the progeny of a cross between two candidates carries only desirable alleles. In a simulated introgression scenario, selecting pairs to cross on the basis of their PCV allowed the alleles to be introgressed into the population faster than truncation selection on GEBVs (Han et al. 2017).

Rather than rank and select individual candidates or specific pairings like OHV and PCV, Kemper et al. (2012)’s method (OHS) chooses a population of candidates that complement each other (that is, a group with optimum “ultimate genotype” GEBV). Goiffon et al. (2017) compared OHS to other selection methods in simulation, including their own derived method, OPV (Optimum population value). OPV selects a set of candidates whose pool of haplotype segments can be stacked to produce the doubled haploid with the highest GEBV. That is, OPV discards the requirement from OHS that the two haplotypes of each chromosome segment in the “ultimate genotype” must come from different candidates. Over most replications of Goiffon et al. (2017)’s simulation experiment, the largest genetic improvement after 10 generations of breeding came from populations under selection on OPV, followed by populations under OHS. Both outperformed selection on OHV or GEBV. Goiffon et al. did not, however, test the performance of OHS or OPV in recurrent applications or over a long period of time. The decisions of OHS and OPV are nontrivial optimisation problems. These studies implemented them using evolutionary computing solvers, rather than an exact solver like the operations research approach of Han et al. (2017).

Here, we investigate the performance of OHS, as implemented in Kemper et al. (2012), when applied over successive generations in a closed wheat population, and across a wide variety of parameters including number of parents available for crossing and family sizes. The aim was to provide breeders with some insight into when OHS might be useful strategy to accelerate genetic gains and maintain genetic diversity. Recurrent rounds of OHS selection were performed in simulation, starting with real wheat founder genotypes. Performance of OHS was compared to recurrent rounds of truncation selection and OCS, and in some cases OPV. Performance was measured in terms of genetic gain, maintaining genetic diversity, and preserving high-quality haplotype segments in the population. We hypothesised that OHS would outperform the other methods in maintaining useful diversity, and therefore, in a closed population, given long time horizons to allow for stacking to occur, could produce larger genetic gains. We also hypothesised that using OHS to select founder lines would provide a good foundation of useful standing variation to be exploited in a subsequent breeding program.

## 2 Methods

### 2.1 Base population

LongReach Plant Breeders provided phased genotypes for a set of 16,565 diverse elite wheat germplasm lines. The genotypes included 5,112 SNPs across 21 chromosomes (with a total genome length of 2,994 centimorgans). A set of additive effects for the 5,112 SNPs, describing grain yield, were also provided. Those SNP yield effects originated from best linear unbiased estimations (BLUEs) on data from 186 field trials between 2012 and 2018. For the simulations in this paper, these were assumed to be true SNP effect values that stayed constant over all generations. All GEBVs discussed in this study are based on this set of marker effects. Hence, the ‘GEBVs’ in the simulations function as true breeding values.

Two base populations of 50 lines were selected from the diverse germplasm. One set (GS-selected founders, abbreviated GS_f_) were selected using truncation selection – i.e., the 50 genotypes with the highest GEBVs. The other set (abbreviated OHS_f_) were selected using OHS on the population of diverse germplasm. Genomic relationships among the base populations are shown in Figure 1. Simulation experiments were carried out using both base populations, and differences in outcomes based on the choice of base population are discussed.

**Fig. 1:**
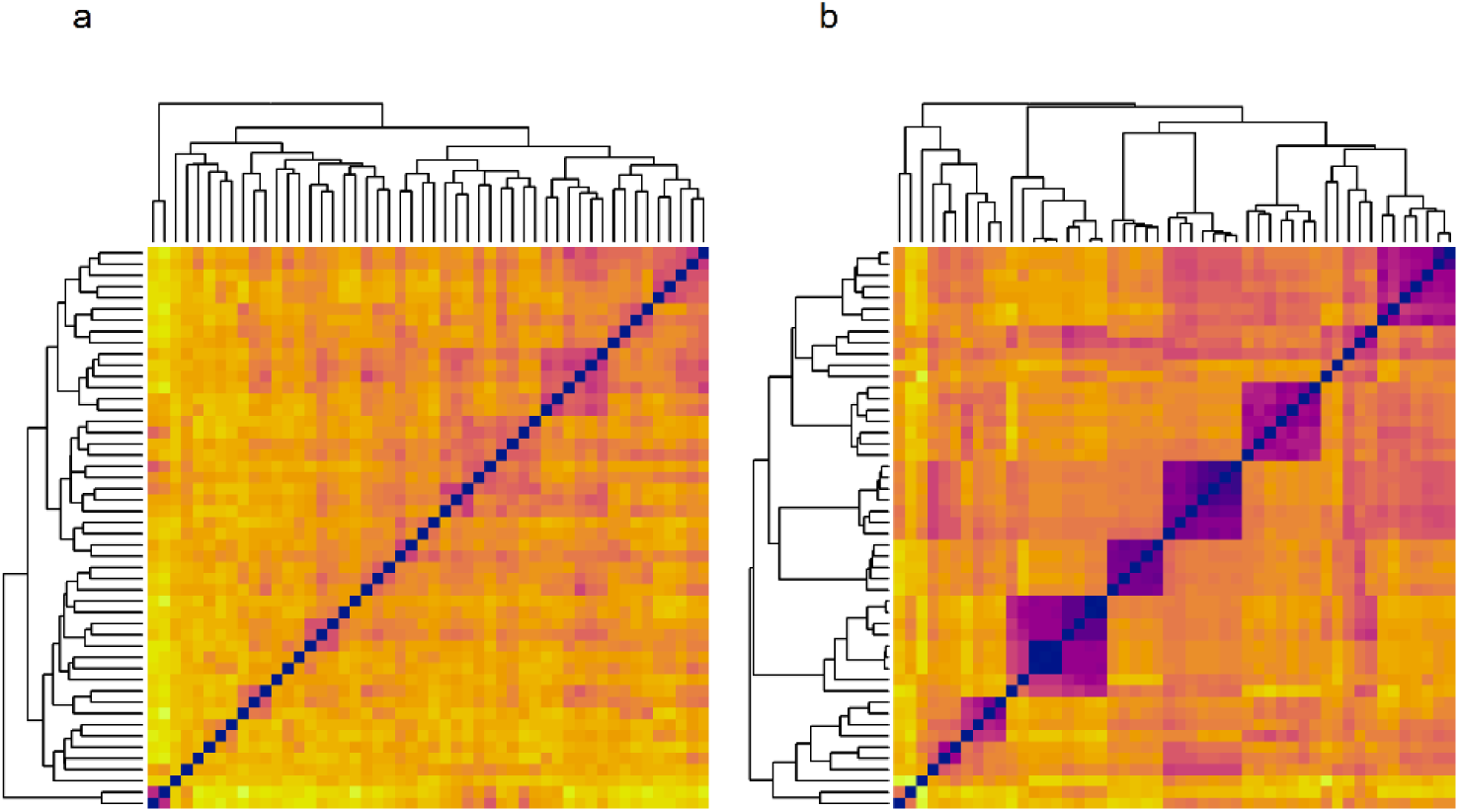
Summary of the base population wheat genotypes. (a) Heatmap of Roger’s genetic distance between the genotypes of the OHS-selected founding population (that is, the 50 candidates selected by Kemper et al. (2012)’s genetic algorithm (GA) from a large diverse population). Dark colours indicate a low genetic distance between pairs of candidates. (b) Corresponding heatmap for genetic distance between GS-selected founding genotypes (that is, the 50 candidates selected by truncation selection on GEBVs from the large diverse population).

### 2.2 Optimal Haplotype Selection

If a GEBV is a whole-genome score representing the estimated contribution of QTLs near to a genetic marker to the phenotype of a candidate organism, we can define a corresponding “local GEBV” that is the estimated contribution of QTLs in a particular genomic region of one haplotype of that candidate:

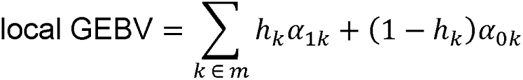

Where *m* is the set of SNP markers belonging to the segment haplotype for which the local GEBV is calculated, *h_k_* is the allele of one SNP marker (either 0 or 1), and *α_0k_* and *α_1k_* are the additive marker effects of alleles 0 and 1, respectively, on marker *k*.

We refer to “haploblocks” as a set of non-overlapping regions of contiguous markers that cover the genome. In the population, there can be up to 2*^m^* unique haplotypes per haploblock (for a haploblock spanning *m* SNP markers). The observed number of unique haplotypes depends on the extent of linkage disequilibrium. Diploid individuals have two haplotypes (and two local GEBVs) per haploblock. The sum of the pair of local GEBVs of all haploblocks adds up to the candidate’s GEBVs.

The goal of OHS is to maximise the score of the “ultimate genotype” that can be constructed from the best haplotypes in the selected population, at each haploblock. Kemper et al. (2012) imposed the restriction that the two haplotypes of each haploblock in the “ultimate genotype” should originate in separate members of the selected population. This restriction is not present in Goiffon et al. (2017)’s corresponding method OPV.

To select using OHS, a user chooses the number *n* of candidates they wish to select out of their candidate pool, and provides the pair of local GEBVs for the user’s chosen haploblocks of each candidate in the pool. OHS will advise the user which *n* candidates should be selected to create a subpopulation with the highest stacking potential (highest-scoring ultimate genotype). Formally, the function that OHS optimises is:

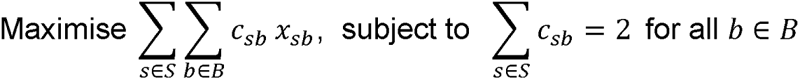

where *S* is the *n*-member subset of the candidate pool that is being optimised; *B* is the set of haploblocks on which the potential founders are scored; *x_sb_* is the higher of the candidate *s*’s two local GEBVs for haploblock *b*; and *c_sb_* are a collection of binary (0 or 1) decision variables that are constrained to select exactly two contributor candidates from *S* for each marker block *b*, out of which the ultimate genotype of the candidate subset *S* is constructed. After preliminary investigations, the number of haploblocks used in the presented OHS simulations was fixed at 105 (each of the 21 wheat chromosomes were dived into 5). The effect of haploblock definition is discussed in the Discussion section.

The *x_sb_* term being the local GEBV for the more favourable of a candidate’s two haplotypes for the haploblock ensures the two haplotypes for that block in the ultimate genotype must come from two different candidates. This does not require that the two haplotype blocks are distinct and the optimal genotype be heterozygous, but does encourage the choice of more candidates that should capture more diversity. It also avoids the possible scenario where one very elite candidate is optimal, so contributes all its haplotypes to the ‘ultimate genotype’, resulting in the rest of the selected population being random choices.

In plant breeding where inbred or doubled haploid lines are the goal, the choice of two haplotypes may seem unnecessary or even counterproductive. Goiffon et al. (2017)’s OPV function is a modification of the OHS/Kemper et al. (2012) fitness function to remove the restriction, meaning it selects a set of candidates for the ultimate fully-homozygous stack. Comparisons between the performance of OPV and OHS are presented in the results.

Choosing a subset that contains the segments to produce the highest-scoring ultimate genotype is a highly combinatorial problem. The number of possible subsets grows rapidly with the size of the population and the size of the desired subset, such that it is completely infeasible to explore all possible subsets to find the optimum. The task of choosing 50 founders from a base breeding population of 200 has over 10^47^ possible subsets of 50 to consider. Therefore, Kemper et al. (2012) used an evolutionary search strategy to produce a ‘good enough’ solution in a defined number of iterations. This study used the genetic algorithm (GA) implementation of OHS produced by Kemper et al. (2012). Further information on the GA implementation’s design is available in Kemper et al. (2012)’s Supplementary files.

Preliminary trials showed that running Kemper et al. (2012)’s GA with an internal population size of 1000 chromosomes (evaluating 1000 subsets *S* of candidates per generation) for 2500 generations of iteration reliably led to acceptable convergence of the fitness function. These hyperparameter values were used when collecting all remaining OHS results in this paper.

### 2.3 Simulation Design

Design of the stochastic simulations in this investigation did not aim to mimic the design of a real-world breeding program, but rather to characterise and give insights into the situations in which OHS might be a useful strategy. To do this, OHS was compared to three other selection strategies: truncation selection on true breeding values, OCS, and OPV. Simulations involved recurrent selection over a long (100 non-overlapping generations) time frame, to test the long-term limits of the selection strategy. The population at each generation were F1 crosses of the previous generation; that is, no inbreeding generations (selfing, or producing doubled haploids) occurred in simulation, and candidates were not inbreds.

The tool genomicSimulation (Villiers et al. 2022) was used to simulate the genotypes of offspring of crosses, as well as to calculate the GEBVs and local GEBVs on which selection occurred. The tool assumes crossovers in meiosis occur with uniform probability across the length of each chromosome, and mutation is not simulated. It was necessary that the simulation tool track the haplotype phase of the genotype, because calculating local GEBVs for OHS requires this information. This is especially relevant in a population of highly heterozygous candidates, like those produced by the recurrent selection without selfing procedure in the simulations, where the two alleles for a marker are often different.

A population size *n* (*n* = 2, 10, 25, 50 or 100) was maintained at every generation of simulation (Figure 2). To progress to the next generation, the population had to be replaced with a population made up of its progeny. Genotypes of F1 progeny from the half diallel of *n* (*n* – 1)/2 crosses between pairs of candidates were simulated. Different simulations tested different numbers of progeny produced from each crossed pair in the half diallel (*p* = 1, 10 or 100). genomicSimulation was then used to produce the table of local GEBVs of the *pn* (*n* – 1)/2 candidates in the offspring pool. The local GEBVs were based on the 105 chromosome segments containing the markers within each 1/5^th^ of the length of each chromosome, as previously described, and were calculated assuming genotypes and marker effects were known with perfect accuracy. Novel haplotypes and local GEBVs could appear between generations, even though marker effects and chromosome segments blocks were fixed, because genomicSimulation’s meiosis simulation allowed some recombination within blocks to Kemper et al. (2012)’s GA, which implements OHS. The *n* candidates from the offspring as well as between blocks. The table of local GEBVs of the offspring pool were then passed pool selected by OHS then became the parents of the next generation (Figure 2b).

**Fig. 2:**
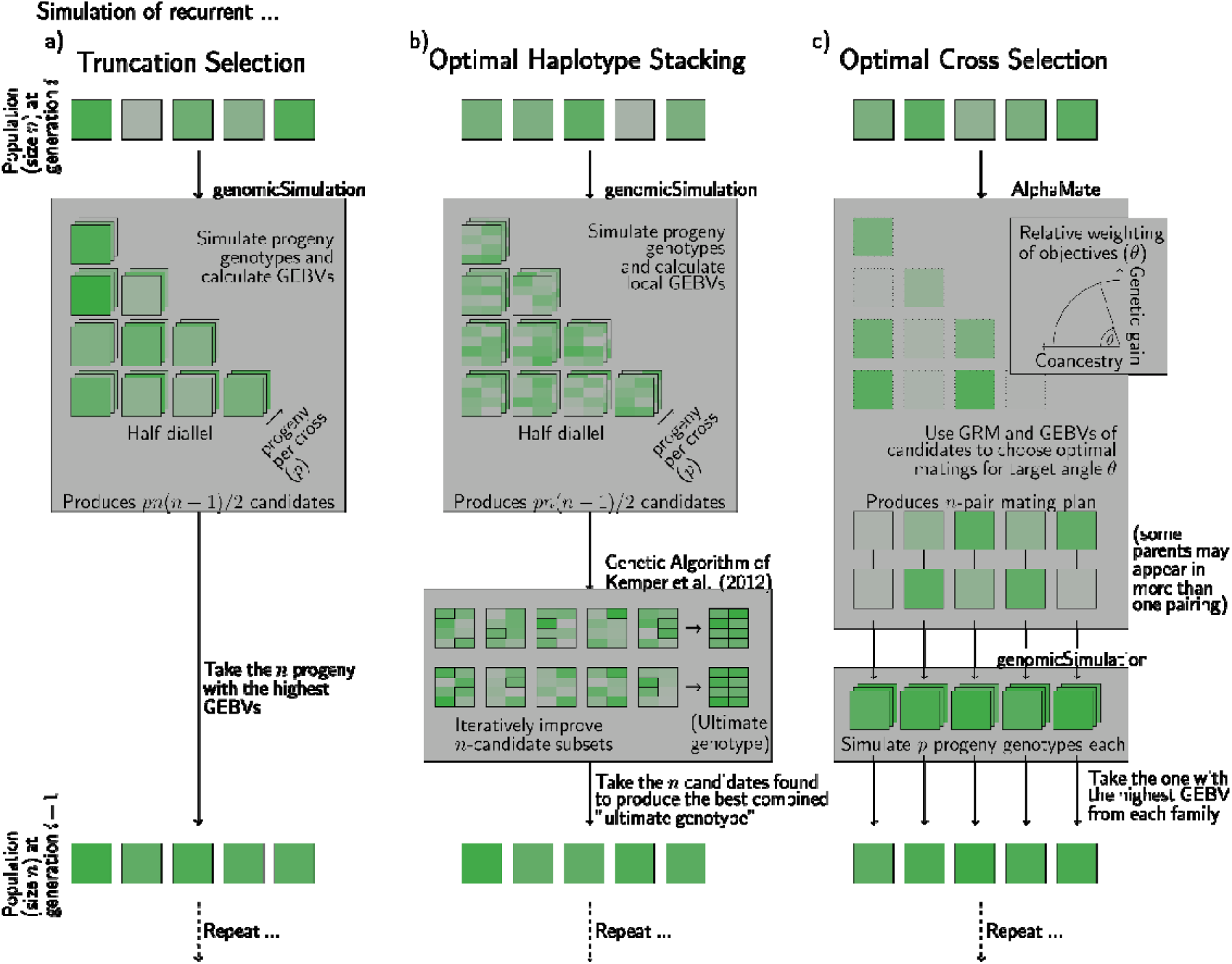
Structure for the three compared simulation experiments. (a) The recurrent truncation selection simulations involve simulating the genotypes and GEBVs of full-sibling progeny of each possible cross between pairs of parent genotypes using genomicSimulation (Villiers et al. 2022), followed by selecting the with the highest GEBVs from the resulting pool of offspring and installing those candidates as the parents of the next generation. (b) OHS (optimal haplotype stacking) simulation structure is initially the same as the truncation selection procedure (simulating full-sibling families for each pair in the half-diallel), but instead of selecting on whole-genome GEBVs, selection is done based on local GEBVs using Kemper et al. (2012)’s genetic algorithm (GA) to find a set of candidates in the offspring pool that contain haplotype blocks that can stack to the best “ultimate genotype”. (c) OCS (optimal cross selection) simulation structure involves using AlphaMate (Gorjanc and Hickey 2018) to choose mate allocations from the set of founder genotypes that would best suit a strategy that balances genetic gain and efforts to minimise coancestry according to a target angle θ between the two objectives. The next generation is then produced by simulating the genotype of full-sibling progeny from each of the crosses in the mating plan, using genomicSimulation, followed by selecting the best one from each family to carry forwards to the next generation. Across all selection conditions, the selected candidates of each selection cycle become the parents of the next generation, so that simulation tests the performance of the different methods over long-term recurrent selection.

Three other selection methods were simulated to compare to OHS. In simulations of truncation selection (Figure 2a), the same process to create an offspring pool of *pn* (*n* - 1)/2 candidates was used. genomicSimulation was then used to calculate the GEBVs of those candidates, based on the ‘true’ additive marker effect set provided by LongReach Plant Breeders, and the candidates with the highest GEBV were selected to become the parents of the next generation.

At each generation in the simulations of OCS (Optimal Cross Selection, Figure 2c), the genomic relationship matrix (GRM) of the *n* candidates was generated, and their GEBVs calculated by genomicSimulation from the unchanging ‘true’ marker effects, as described above. The tool AlphaMate (Gorjanc and Hickey 2018) was used to select optimal matings from those *n* parent-candidates. genomicSimulation was then used to simulate genotypes of the offspring resulting from the AlphaMate mating plan. Like in the other selection conditions, several simulation conditions using different values for the family size/number of progeny per cross (*p*) from each cross in the mating plan were carried out. The candidate with the highest GEBV in each resulting family was then selected to be one of the *n* parents of the next generation. This design for OCS using the same parameters *n* and *p* as the truncation selection and OHS conditions allowed for straightforward comparison of results. Results from this particular OCS program design are presented because other recurrent OCS-based simulation designs tested (involving selecting from a half-diallel offspring pool, or involving matching its coancestry or genetic gain objectives to the coancestry or genetic gain measures recorded in other simulation conditions) showed much lower rates of gain and poorer performance. AlphaMate requires a target angle parameter to define the relative importance of minimising coancestry and maximising genetic gain in its objective function. For each combination of parameters *n* and *p*, grid search with a coarseness of 5 degrees, followed by binary search to a coarseness of 1 degree, was used to identify the target angle which gave the highest long term gain (that is, the highest average GEBV at generation 100). Only the results from running AlphaMate with this target angle are presented.

Finally, simulations were run that used the same design as OHS simulations (Figure 2b) but replacing the OHS fitness function with a near-identical fitness function that only selected one haplotype per chromosome segment (as in Goiffon et al. (2017)’s OPV).

### 2.4 Summary and analysis

Simulation of each experimental scenario was run independently 20 times for all but the largest OHS scenarios (100 progeny per cross, and a population size of 50 or 100), which, due to runtime issues, were only replicated 10 times. Four summary statistics were calculated for the population at each generation of each simulation trial: mean GEBV, upper selection limit (the GEBV of the ultimate candidate that could be constructed at the current generation, Cole & VanRaden 2011), mean coancestry, and a gene diversity index from Nei (1973). The results show these summary measures averaged across replications.

The first summary statistic, the mean GEBV, is the mean of the breeding values of all the genotypes in the population at a given generation. Breeding values at all generations in simulation are calculated using a ‘true’, consistent set of marker effects, as previously described.

The second is Cole & VanRaden (2011)’s upper selection limit: the GEBV of the genotype created by taking the best allele present in the population at every marker. Since the marker effects in this study are purely additive, that genotype will be fully homozygous. Assuming no introduction of fresh diversity, this is the maximum GEBV that descendants of this population could achieve. Unlike the stack-of-segments ultimate genotype optimised in OHS, the upper selection limit is based on the best possible choices at the marker level, so may include chromosome segments that do not exist yet.

The third summary statistic, the mean coancestry of the population at a given generation, is calculated as twice the mean of the values in the genomic relationship matrix, including the diagonal, with the reference population (base) being the founder set, OHS_f_ or GS_f_ (Gorjanc and Hickey 2018; Isik, Holland, and Maltecca 2017).

Finally, the last statistic was the mean of Nei (1973)’s gene diversity index *H_e_* among a given generation’s population, calculated as follows:

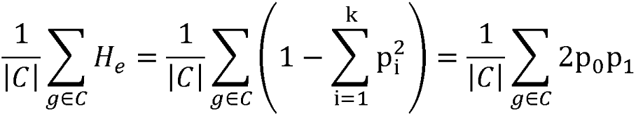

where *g* is a candidate in *C*, the breeding population at a given generation. |*C*| is the size of *C*, or the number of candidates in the population, and p_0_ and p_1_ are the proportions of the alleles of that candidate that are (respectively) 0 and 1. The simplification is possible thanks to the assumption of Hardy–Weinberg equilibrium, and all markers being SNPs (thus *k* = 2) that have been re-coded so that their reference and alternate allele are 0 and 1 regardless of what nucleotides the actual alleles are.

## 3 Results

### 3.1 Truncation selection versus OHS for genetic gain and maintaining genetic diversity

The base population selected by OHS was considerably more diverse than the base population selected by truncation on GEBVs (Figures 1a and 1b). The same pattern is seen in Figure 3: OHS selection samples more evenly from the available genetic diversity than truncation selection on GEBVs, and conversely, truncation selection strongly favours breeding offspring from only a few founders. This effect is more obvious when the population is more diverse (Figure 3a). Initially, recurrent truncation selection achieves higher rates of gain than OHS, but rates of gain then fall to zero as diversity is exhausted (Figure 4). Meanwhile, OHS produces lower but steadier rates of gain. After 100 generations of intercrossing in a closed population (Figure 4), the rate of gain with OHS has only slightly decreased, and the overall gains in yield GEBV values are higher than those from recurrent truncation selection (which has reached a plateau). For the condition pictured in Figure 4 (10 progeny per cross, and selecting 25 genotypes each generation), the OHS approach begins to outperform the truncation strategy on the same base population after 70 generations.

**Fig. 3:**
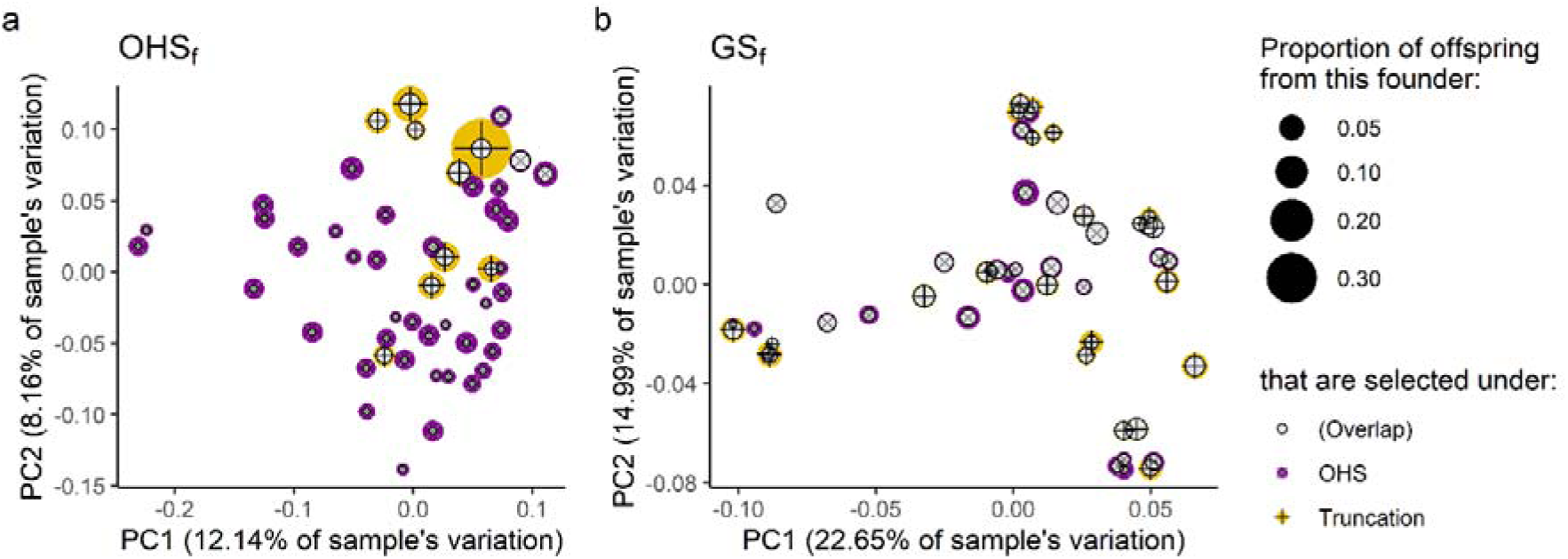
Diversity of founder population and proportion of lines selected by truncation and optimal haplotype stacking (OHS) selection. (a) Principal component plot of OHS_f_, the founding population of 50 lines selected by OHS from a large diverse population. The size of the scatter points corresponds to the proportion of the selected progeny that have that founder as a parent. Proportions of first generation offspring selected by OHS and of first generation offspring selected by truncation selection are shown in different colours. (b) Corresponding principal component plot of GS_f_, the founding population of 50 lines selected by truncation selection from the large diverse population.

**Fig. 4:**
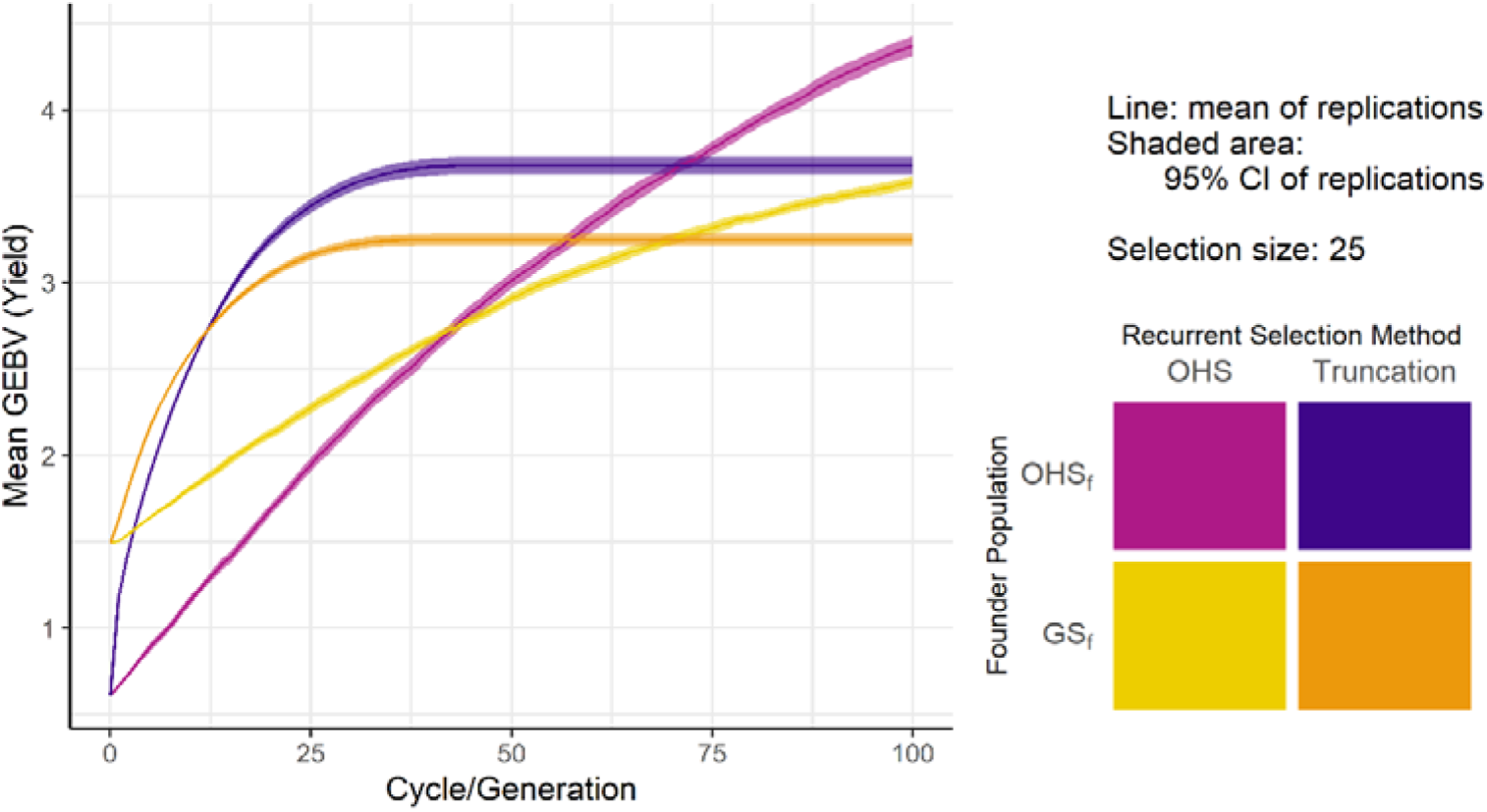
Results of simulating recurrent truncation and OHS selection in a wheat breeding program using different founder populations. Plot shows each generation’s mean and 95% confidence interval of GEBV, comparing simulations starting with OHS-selected (OHS_f_) and truncation-selected (GS_f_) founder sets and running 100 generations of OHS recurrent selection or recurrent truncation selection. The simulation runs in this plot use the parameters 10 progeny per cross and recurrent selection size of 25.

The plateau in genetic gain observed in recurrent truncation selection conditions corresponds to the point where genetic diversity in the population is exhausted (Figure 5a and 5b). Average GEBV in OHS simulation runs can surpass average GEBVs from truncation selection conditions in the later generations, because OHS maintains more genetic diversity (Figure 5b) and more useful genetic diversity (Figure 5c), where “useful” diversity is defined as haplotypes with the highest GEBV in their haploblock. Even after 100 generations, this stock of genetic diversity is not exhausted, so recurrent OHS could continue seeing gains beyond 100 generations.

**Fig. 5:**
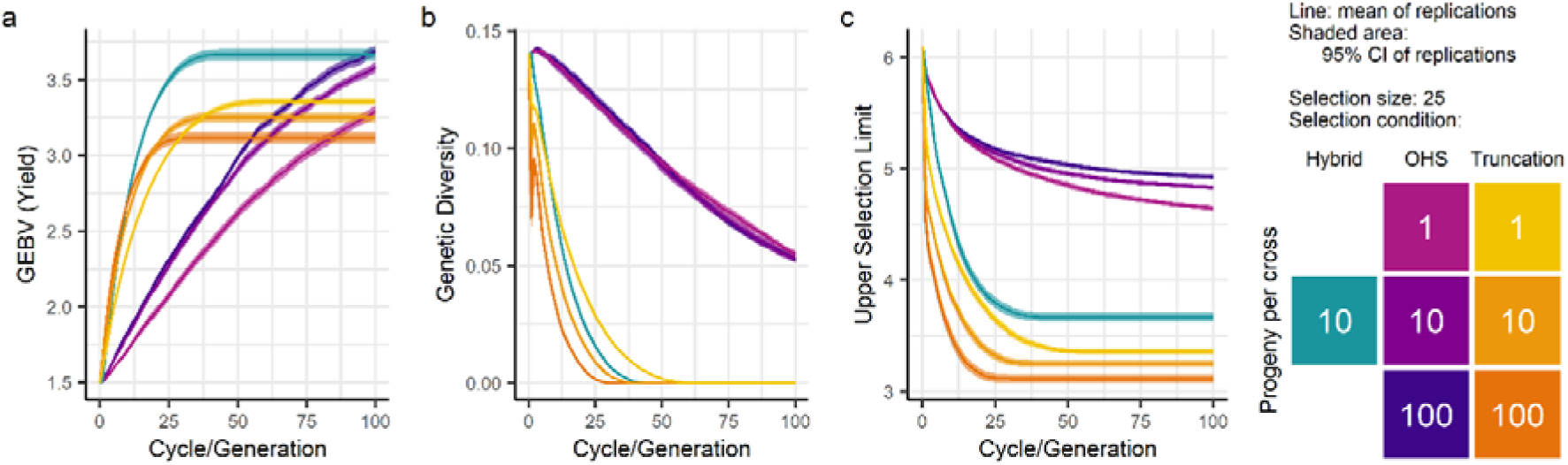
Comparison of truncation selection and OHS in recurrent selection in a closed population based on three performance criteria. (a) Average GEBV of the population over generations. (b) Genetic diversity over generations (Nei (1973)’s expected heterozygosity measure). (c) Upper selection limit (Cole & VanRaden 2011) at each generation. Results are from simulation with a recurrent selection size of 25, with founder population GS_f_. Represented in the plots are the following conditions: selection by OHS with 1, 10 or 100 progeny per cross; truncation selection with 1, 10 or 100 progeny per cross; and a ‘hybrid’ condition with 10 progeny per cross, where selection is done by OHS in the first generation and by truncation on GEBVs for every generation thereafter.

The ‘hybrid’ selection strategy in Figure 5 is simulated by performing selection using OHS in generation 1, followed by recurrent truncation selection for the remaining generations. This strategy’s genetic gains are higher than those of the corresponding truncation selection condition after 10 generations, even with one generation of slow OHS gains. While its genetic gains plateau like the other truncation selection conditions, the average GEBV at the hybrid condition’s plateau is notably higher than corresponding pure truncation selection conditions. The gains from recurrent truncation selection on the OHSf base population surpasses recurrent truncation selection on the GS_f_ base population (Figure 4) in a similar time frame, and also achieves a notably higher plateau. The ‘hybrid’ selection approach therefore surpasses pure recurrent truncation selection in both the long-term and shorter term. Compared to the truncation condition with the same parameters, the hybrid condition’s useful diversity is slightly higher (Figure 5c) and it exhausts diversity a few generations later (Figure 5a-b). Other simulated ‘hybrid’ conditions show that the long-term improvement in genetic value is smaller if the OHS generation is later than generation 1, suggesting the extent of the benefit depends on the amount of diversity in the population.

OHS selection benefits from more progeny per cross (Figure 5, 6, 7). In early generations, truncation selection conditions with more progeny per cross likewise show higher GEBVs (Figure 5a, Figure 6). However, in later generations, after the plateau is reached, truncation selection conditions with more progeny per cross have lower GEBVs than their few-progeny counterparts (Figure 5a, Figure 7).

**Fig. 6:**
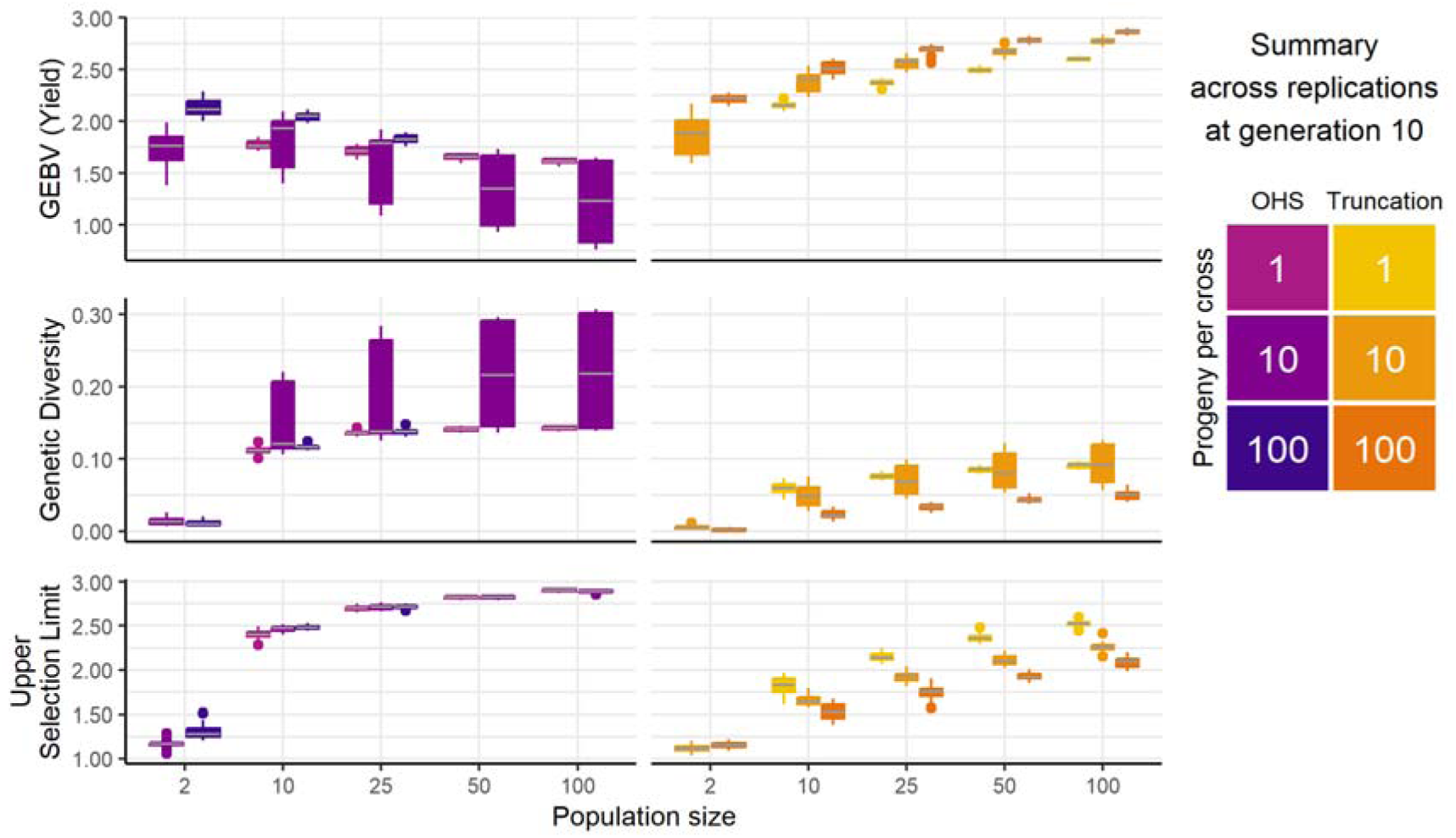
Comparison of optimal haplotype stacking (OHS) and truncation selection after 10 generations, across all combinations of population size and progeny per cross parameters tested in simulation.

**Fig. 7:**
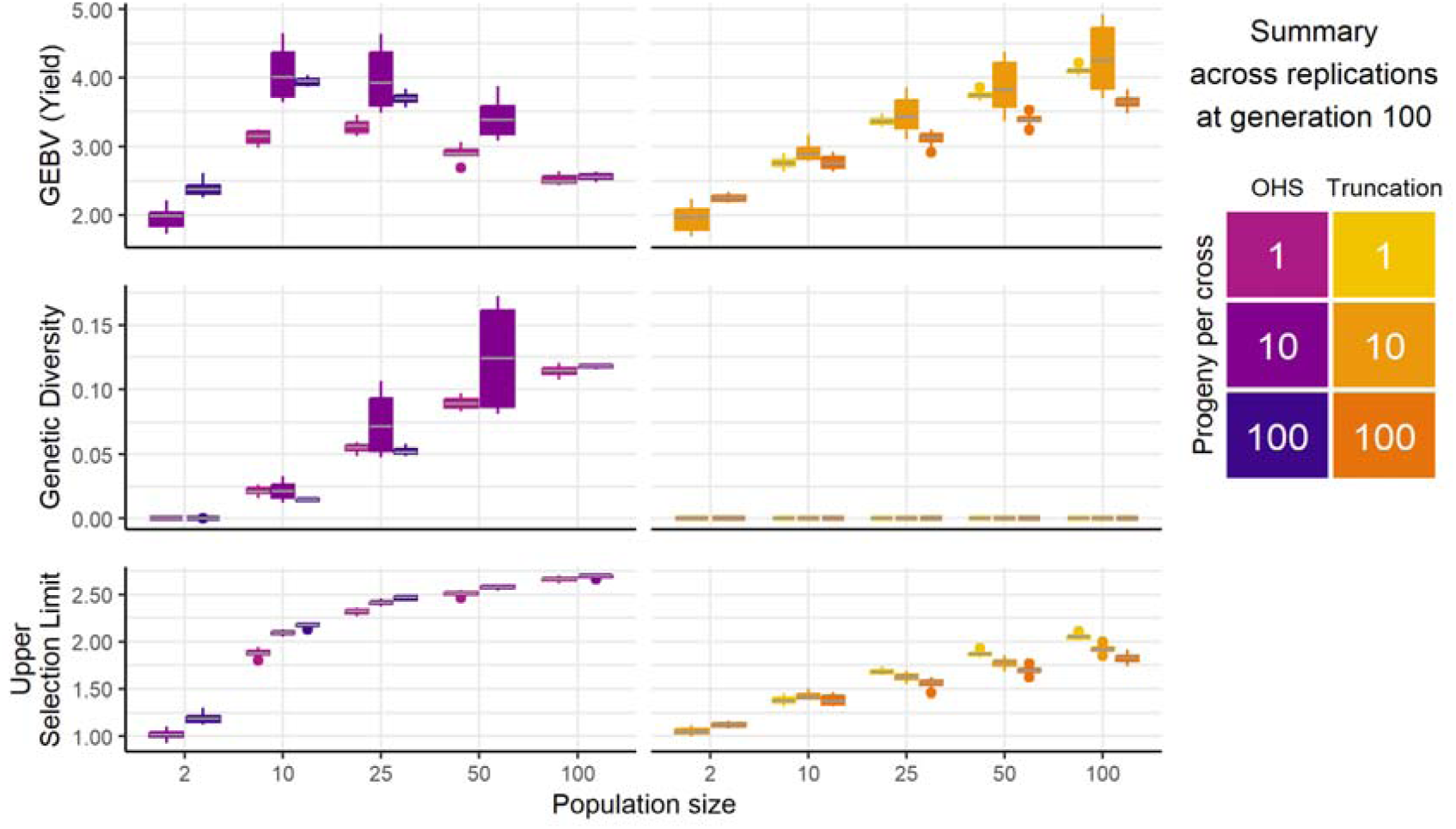
Comparison of optimal haplotype stacking (OHS) and truncation selection after 100 generations, across all combinations of population size and progeny per cross tested in simulation. For all truncation selection scenarios, genetic diversity was exhausted (0) by generation 100.

While truncation selection benefits from higher population sizes for achieving higher rates of genetic gain, the performance of OHS selection relative to truncation selection improves with lower population sizes (Figure 6, Figure 7). When the population size is 2, both selection conditions predictably exhaust their diversity too fast to see much genetic gain. However, average GEBVs from OHS selection are higher when the size of the selection is 10 than when it is 25, and higher when it is 25 than when it is 50 or 100. Larger population sizes, however, allow OHS to maintain more diversity than smaller ones. By generation 100 (Figure 7), truncation selection conditions had exhausted all genetic diversity under Nei (1973)’s index, while all OHS conditions with population sizes greater than 2 maintained diversity and had higher upper selection limits than the truncation selection plateau.

### 3.2 Optimum cross selection versus OHS for genetic gain and maintaining genetic diversity

The best recurrent OCS programs simulated outperformed truncation selection on genetic gain both in the relatively short-term and in the long-term. Within 10 to 25 generations, OCS produced higher average GEBVs than truncation selection (Figure 8a). The OCS conditions take 50 to 75 generations to exhaust diversity, while truncation selection conditions exhaust theirs before 40 generations (Figure 8d). Because the OCS objectives in the pictured OCS strategies were highly weighted towards genetic gain (Table 1), the coancestry of pictured OCS strategies does not lag far behind that of truncation selection strategies (Figure 8b). The OCS strategies considered maintain a higher upper selection limit than truncation selection, though the trend has the same profile as truncation selection and is distinctly lower than the upper selection limit maintained by OHS (Figure 8c). Larger family sizes in OCS produced faster GEBV gains and slightly higher GEBV plateaus (Figure 8a) for the same rate of coancestry increase (Figure 8b). Compared to these, OHS had lower and steadier rates of gain, gained coancestry slower, and maintained more diversity, even after 100 generations.

**Fig. 8:**
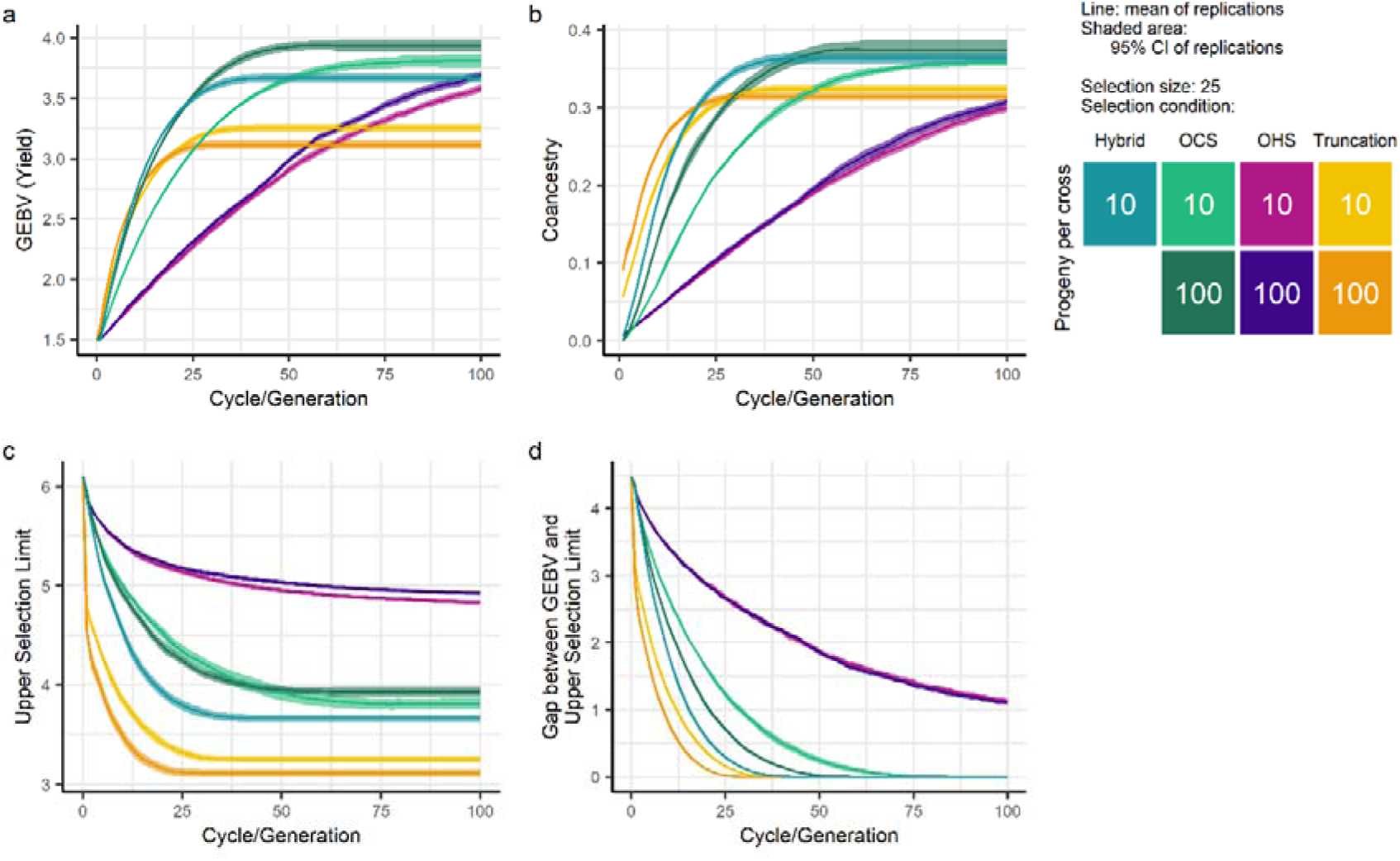
Simulation results comparing optimal haplotype stacking (OHS), truncation selection, and optimal cross selection (OCS). (a) Trends in each generation’s mean GEBV. (b) Trends in each generation’s genetic diversity (Nei (1973)’s expected heterozygosity measure). (c) Trends in each generation’s upper selection limit. (d) Gap between each generation’s mean GEBV and upper selection limit. Results are from simulation with a recurrent selection size of 25. Represented in the plots are the following conditions: selection by OHS with 100 progeny per cross; truncation selection with 10 progeny per cross; a hybrid selection condition involving one generation of OHS followed by 99 generations of truncation selection, with 10 progeny per cross; OCS where the crossing plan designed the same number of offspring as are present in the half diallel for truncation and OCS conditions; OCS involving producing 10 progeny from each pairing in the crossing plan, and selecting the best of each of those families by GEBV; and OCS involving producing 100 progeny from each pairing in the crossing plan and selecting the best of each of those families by GEBV.

**Table 1:** Parameter combinations tested in simulation experiments. Each optimal haplotype stacking (OHS), truncation selection, or optimal cross selection (OCS) condition can be simulated with a population size of 2, 10, 25, 50 or 100, and 1, 10, or 100 full-sibling progeny per cross (“ppc”). Conditions with population size 2, and 1 progeny per cross, were not simulated for truncation selection or OHS: there is only one possible pairing in the half-diallel of a population with 2 members, so producing only one progeny per cross at that population size is below the replacement rate needed for the next generation. OCS uses a target angle parameter to define how to weight the two objectives of increasing genetic gain and minimising coancestry gain. Grid search with a coarseness of 5 degrees, followed by binary search to a coarseness of 1 degree, was used to find the target degree that maximised long term genetic gain in simulation (highest population GEBV average after 100 generations). The identified target angles are shown in the table.

|  |  | Select 2 | Select 10 | Select 25 | Select 50 | Select 100 |
| --- | --- | --- | --- | --- | --- | --- |
| OHS | 1 ppc |  | X | X | X | X |
|  | 10 ppc | X | X | X | X | X |
|  | 100 ppc | X | X | X |  |  |
| Truncation | 1 ppc |  | X | X | X | X |
|  | 10 ppc | X | X | X | X | X |
|  | 100 ppc | X | X | X | X | X |
| OCS | 1 ppc |  | 58° | 68° | 70° | 73° |
|  | 10 ppc |  | 85° | 86° | 86° | 88° |
|  | 100 ppc |  | 87° | 86° | 88° | 89° |

Interestingly, the ‘hybrid’ selection condition (where one generation of OHS was followed by recurrent truncation selection for the remaining generations) outperformed the OCS condition with the same number of progeny per cross in the short-term, producing the similar rates of gain to the OCS condition with an order of magnitude more progeny per cross. However, the hybrid selection condition’s upper selection limit dropped faster than the OCS conditions’, and it exhausted its diversity at the same point as truncation selection, so it had a slightly lower GEBV plateau than the OCS conditions.

### 3.3 Targeting one haplotype versus targeting two haplotypes per chromosome segment for genetic gain and maintaining genetic diversity

The value and trend of the upper selection limit (Figure 9c) was similar for recurrent application of OPV and OHS. As expected, the genetic diversity of the population (Figure 9b) is higher under OHS, which imposes the constraint that each pair of haplotypes in the ultimate genotype must come from two separate candidates, than it is when this criteria is relaxed (OPV). Interestingly, the genetic gain from OHS is higher than the genetic gain from OPV in our simulation results (Figure 9a).

**Fig. 9:**
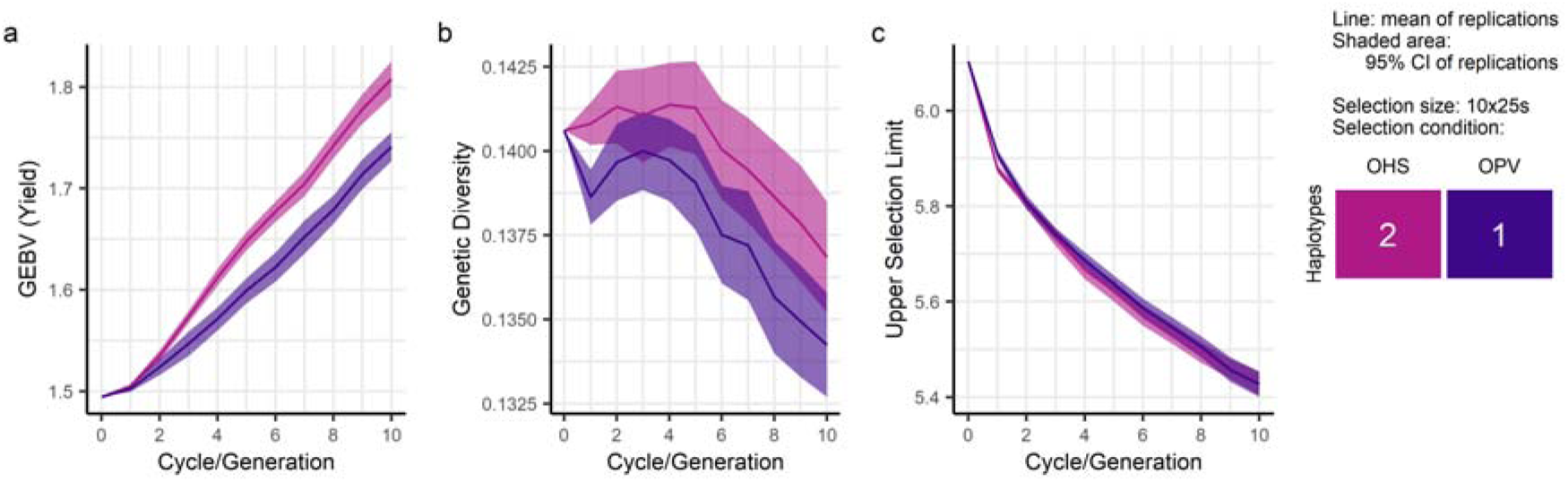
Simulation results for 10 generations of optimal haplotype stacking (OHS) compared to Goiffon et al. (2017)’s optimal population value (OPV), which uses a near-identical fitness function but selects on the best single haplotype at each chromosome segment where OHS selects on the best two haplotypes (from two different candidates) at each chromosome segment. (a) Trends in each generation’s mean GEBV. (b) Trends in each generation’s genetic diversity (Nei (1973)’s expected heterozygosity measure). (c) Trends in each generation’s upper selection limit. Results are from simulation with a recurrent selection size of 25, and 10 progeny per cross. 10 generations of simulation results are shown, but these trends continue when simulated out to 20 generations.

## 4 Discussion

### 4.1 OHS is a selection method that maintains genetic variation and produces steady genetic gain

Our simulation results demonstrate that OHS as a recurrent selection method can result in greater long-term gains than either recurrent truncation selection or the OCS program designs simulated. OHS shows the greatest gains over truncation selection when small populations of parents are selected and mated each year with large family sizes. For large population sizes, truncation selection was often superior to OHS in terms of genetic gain after 100 generations of simulation.

OHS may be less desirable than OCS or truncation selection in practical breeding programs because while it retains potential for impressive improvements, it does not produce these improvements very fast. It takes 25 generations for OHS to outperform truncation selection for a very small breeding population of 10 individuals, and the time to exceed performance of other strategies is longer as the breeding population size increases: Figure 5a shows OHS takes 60 generations to outperform truncation selection when the breeding population is 25 individuals. In contrast, the best OCS condition outperforms truncation selection at that population size after only around 10 generations (Figure 8a).

A distinguishing feature of OHS in our results was how much genetic diversity was maintained compared to other selection methods. Truncation selection and OCS conditions saw a sharp, exponential initial decline in diversity, while the trend of OHS conditions’ genetic diversity measures was a much shallower, more linear decrease. This may be because OHS can select some mediocre candidate genotypes which would not be selected by truncation selection, because they have a few good haplotypes. The other, mediocre, haplotypes of such candidates will likely be different to the higher-value haplotypes of other selected candidates, which would add to measurements of genetic diversity, without necessarily meaning that the increased diversity is valuable for breeders. However, logging the upper selection limit showed that OHS does not simply maintain more genetic diversity than other methods, but maintains more high-value haplotypes.

### 4.2 Parameters for the highest rates of gain with OHS

The relative performance of OHS improved in both the long and short term with higher numbers of progeny per cross. Full- or half-siblings, no matter how high their GEBVs, are likely to get that GEBV from many of the same high-scoring segments, and therefore OHS’s fitness function will not select them together. Higher numbers of progeny per cross do however mean greater chances of a desired recombination between favourable alleles occurring, so more chance of there being a superior recombinant to select from the pool.

Improvements in performance from more progeny per cross in OHS are diminishing, with a tenfold increase from 1 to 10 progeny per cross improving performance more than the second tenfold increase from 10 to 100 progeny per cross (Figure 5a). A balance between OHS performance improvement and the increased genotyping costs and diminishing returns of more offspring per cross must exist. But overall, this suggests that OHS would be most powerful in species where affordable genotyping platforms exist and large numbers of progeny of the same cross can be produced and selected between.

Larger breeding populations mean it is possible to retain a wider variety of genotypes with more diverse and useful alleles. Both truncation selection and OCS perform relatively better with larger population sizes, because they will maintain more diversity to support future gains. However, the genetic gain achieved by OHS over a given time horizon was relatively higher for smaller populations. For example, our simulations with 10 or 25 parents each generation (Figure 7). In a bottleneck scenario, the ultimate genotype’s segments must come from a limited number of candidates. This increases the likelihood that recombination events make progress towards the ultimate genotype, so OHS on small populations will observe higher rates of gain. Given instead a large selection size, OHS can pick exceptional segments isolated in otherwise low-value genotypes. This then requires more rare recombination events to stack the segments before substantial genetic progress towards the ultimate genotype can be observed. This characteristic is visible in Figure 7: the gap between the mean generation-100 GEBV and the score of the upper selection limit is larger for larger population sizes.

The effect of increasing the number of haploblocks for OHS is very similar to the effect of increasing the population size. Having more haploblocks allows OHS to stack more favourable QTL alleles from different origins to create the ultimate genotype, but does result in very slow rates of improvement of the population, whereas fewer, larger, haploblocks allows for faster rates of gain. The genetic merit of the ultimate genotype will be lower with large block sizes if there are multiple QTL per segment. OPV, the method similar to OHS that instead maximises the upper selection limit of the doubled haploid, has the same behaviour: more haploblocks require a longer breeding program in order to approach the creation of the ultimate genotype (Goiffon et al. 2017).

### 4.3 Potential applications of OHS to breeding programs, and other future directions

As presently implemented and parametrised, selection pressure of OHS (and OPV) can only be controlled indirectly. Selection pressure is a function of the number of candidates being selected from the population, and the number of haploblocks into which the genome is divided. A method for choosing values for these parameters to target particular time frames for improvement, or desired rates of gain, has not yet been developed, but could potentially greatly enhance the practical value of haplotype stacking methods.

Currently, marker effects are estimated with the assumption they will be used to calculate a candidate’s whole-genome GEBV. If haplotype stacking selection methods became important, techniques could potentially be developed to model epistasis and G x E interactions differently, at the block-level, so as to be beneficially incorporated into a block-based selection metric.

Combining OHS with other methods could retain its advantages with faster rates of gain. Our simulations showed that ‘hybrid’ strategy, involving a single generation of OHS followed by truncation selection, could outperform the OCS and truncation selection conditions with corresponding population sizes and numbers of progeny per cross in the short and medium term. However, the hybrid strategy exhausted its diversity a few generations faster than OCS, due to the rapid diversity loss during the recurrent truncation selection phase. Further investigations could combine a single generation of OHS with OCS instead of truncation selection, or interweave OHS generations throughout the breeding program.

Because it identifies useful candidates to retain at the block level, rather than the whole-genome level, OHS could be well-suited to early generations after the introduction of new diversity to a breeding program. When new alleles are carried by individuals with comparatively low overall GEBV, such block-level resolution seems a good tactic to guide selection in integrating new and old alleles. OHS may also perform well in tasks that can be approached with a ‘stacking’ mindset, such as breeding for multiple traits (e.g. stacking disease resistances), breeding for a new complex trait, or targeted exploration of gene bank germplasm resources.

Gorjanc et al. (2016) discuss the problem of ‘masking’ in pre-breeding programs, where desirable chromosome segments (e.g. chromosome segments with favourable effects on the target trait) are contained in the genome of candidates with an overall low GEBV, and so are hidden to whole-genome-score selection measures. The OHS selection method addresses this masking problem by targeting segments as selection units directly, and by selecting at the set-of-candidates level rather than at the individual-candidate level. Candidates can be selected because they have favourable haplotypes not present in other selection candidates, or good haplotypes in haploblocks where the average haplotype value is low, even if the candidate does not have a high GEBV. The authors of Allier et al. (2020) suggested that a stacking method able to find the set of donors most compatible to a set of elite lines, comparable to Han et al. (2017)’s single-donor PCV, would be valuable in pre-breeding programs for enriching the genetic diversity of elite populations. OHS is an indirect stacking method (it does not consider the probability of stacking its selected segments the way that PCV does), but it is used to select sets of candidates that are compatible in a segment-stacking sense. Therefore, OHS matches the approach that Allier et al. (2020) suggested might suit this problem. We did observe, in all scenarios simulated, that OHS retained more “useful” diversity in the breeding program, where “useful” is defined as the haplotype in each haploblock with the highest GEBV.

Selection methods that select specific mating pairs (eg PCV) or specific candidates (eg OHV), rather than selecting the pool of candidates as a whole, lose the ability to select based on what is already represented in the pool, and so lose some of the advantages of OHS and OPV in scenarios around integration of donor and elite lines. Furthermore, in Goiffon et al. (2017), OHV was very frequently outperformed by OHS and OPV over 10 generations of selection. OCS and similar methods (GOCS, Genomic Mating, UCPC) do have the ability to take the composition of a set of candidates into account. It remains to be seen if these methods could, in a simulation design better suited to them or better approximating a practical breeding program, maintain useful diversity to comparable levels to OHS. The ‘boost’ in long-term gains from a single generation of OHS does, however, not seem to be something shared by the OCS family of selection strategies.

We cannot yet judge whether applying OPV would be more appropriate than applying OHS for a breeding program aiming to produce inbreds or doubled haploids. To test this would require different simulated breeding program designs. However, the fact OHS had higher population improvement on average than OPV in this study’s simulations suggests that OHS would be at minimum worth investigating alongside OPV, despite intuitions that OPV would be better suited. OHS maintained more diversity than OPV (Figure 9b), so it may be preferred for breeding goals where diversity is beneficial (eg long-term selection programs, genetic resource exploration or disease resistance breeding). A combined strategy would also be worth investigating, where several generations of OHS were followed by generations of OPV to produce inbred lines.

Although the 100-generation time horizon in our simulation study seems like a long time, speed-breeding approaches (Watson et al. 2018) are increasingly adopted and can turn over up to six generations of spring wheat each year. If the 100 generations of intercrossing, as simulated in this study (assuming fast genotyping and no inbreeding), are performed under speed breeding conditions, this would equate to just 16-17 years. As speed breeding becomes more widespread, breeding programs may need to manage diversity more carefully. OHS, even with its slow rates of gain, could also become increasingly practical as the amount of real-world time it takes for OHS to outperform the other selection methods decreases. However, note that OHS outperforms OCS and truncation selection in our simulations because they exhaust the genetic diversity of the population, and, due to the lack of mutation (or other injection of new diversity into the population) in the simulations, cannot regain it. Whether OHS would outperform them in the long term in real populations would depend on the rate the lost useful diversity can be replaced, by mutation or by management strategies.

For a potential practical application of recurrent OHS, consider the two-part breeding program design proposed by Gaynor et al. (2017). Recurrent OHS (on a population repeatedly inter-crossed and never inbred), as presented in this paper, might suit the population improvement component of a two-part breeding program, as the hybrid condition has shown that the gradual stacking of good haplotypes by OHS provides a good foundation for a faster selection method to achieve higher gains. The component of the breeding program using OHS would then be separate from the product development component of the pipeline, which develops inbreds. This two-part design would permit the use of OHS despite its lower rates of gain, if a product development pipeline that extracts value from the population in a reasonable timeframe could be developed. Alternatively, a possibility is that speed breeding could be used in the population improvement component to compensate for the relatively low rates of gain of recurrent OHS.

### 4.4 Limitations of simulation design

The simulation design – with its isolated, closed population, constant marker effects, true breeding values, perfect genotyping and phasing, and lack of mutation – is not equivalent to a real breeding program.

It was assumed that markers perfectly tracked the QTL over 100 generations of simulation; the controlling QTL had only additive effects; the positions and order on the chromosome of all markers were perfectly known; and for the purposes of calculating local GEBVs for OHS, phase could be perfectly determined. QTL effects could have been simulated, however, the challenge then would be to assume the number of QTL and the distribution of their effects.

In a real-world situation, the estimates of marker effects and estimates of phase could be iteratively updated with phenotype data collected in each new season’s trials. Something else not considered in these simulations is that in practice, the predicted effects of rare haplotype blocks will be regressed hard towards the mean if BLUP methods are used, so the likelihood of identifying a rare and beneficial haplotype block and preserving it in the population might be low. Addressing this might require exploring methods that suggest matings and trial designs intended to increase the frequency of rare haplotypes in the population and their subsequent level of phenotypic recording.

Another limitation of our study is that no mutation was simulated, even though beyond 20 generations of artificial selection, the contribution of mutation to genetic variance becomes quite important (Dudley and Lambert 2004; Barton and Keightley 2002; Keightley 2004). If marker effects that capture the new mutation’s value, or updated SNP arrays, were available, truncation selection and OCS conditions would be able to improve past their plateaus. However, simulating mutation is challenging because the distribution of mutation effects is poorly defined. Future studies will investigate the effect of mutation and erosion of linkage disequilibrium between markers and QTL over time.

One of the assumptions built into the simulation tool was that crossovers occur with uniform frequency along the length of each chromosome. Building into the GA’s objective function some measure of recombination frequency, for example from estimated LD block structures, could improve its suggestions in real organisms.

Usually, in a breeding program, underperforming candidates would be culled before selection, whether by breeders or by failure to survive. This kind of culling was not present in this study’s simulation design. Goiffon et al. (2017) found that OPV often outperformed OHS after 10 generations of selection, but our study’s simulation results had OHS outperforming OPV over the same measure and time-frame (Figure 9a). Goiffon et al. (2017)’s simulations included an extra parameter representing the percentage of the population (the percentage with the lowest GEBV) that would be culled before selection. They found that 70% culling rate for OPV and 80% for OHS produced the highest genetic gains after 10 generations. The difference may be due to different founder populations in the two studies, or it may be that the pre-selection culling of low GEBV candidates conducted in Goiffon et al. (2017) was important to optimise gain under OPV.

Simulation results for a recurrent mate-selection OCS program producing one progeny per (suggested) cross showed that design did not outperform recurrent truncation selection either in short term rates of gain or in long-term increase in genetic merit, for any weighting of the two target objectives. A similar result was observed by Labroo & Rutkoski (2022)’s simulation study, which used one progeny per cross in their OCS condition and found OCS underperformed truncation selection for traits with high heritability. The same scenario, producing multiple progeny per (selected) cross then selecting the best candidate from each resulting family, did surpass recurrent truncation selection within 10-25 generations of recurrent selection in simulation, a result matched by other simulation studies (Allier et al. 2019, Sanchez et al. 2023). Unlike these studies, this study did not incorporate the production of doubled haploids or otherwise remove the heterozygosity from the selected offspring before using them as parents for the next cycle of OCS. Nevertheless, the trends in genetic improvement in these simulated OCS conditions were comparable.

Economic equivalences between different conditions and parameter choices were also not considered during comparisons of simulations.

### 4.5 Conclusion

OHS is a selection strategy which provides steady rates of genetic improvement even over long term horizons, by maintaining useful segments and capturing them as they gradually stack together in progeny of selected crosses. Simulation results have shown that the haplotype stacking approach taken by OHS and the derived method OPV does an excellent job of maintaining the diversity of a closed breeding population, and in the long-term (100 generations) achieves genetic gain comparable to or in some cases exceeding simulated recurrent truncation selection and OCS. The highest rates of gain from OHS, relative to truncation selection, are observed in small breeding populations (10 parents or so) with large family sizes.

In this study’s simulations, OHS outperformed OPV. This result differs from the results observed by Goiffon et al. (2017), but more investigation would be necessary to determine what characteristics of the population and of the breeding program suit OHS better than OPV.

A result from this study’s simulations that may be of interest to breeders is that a single generation of selection with OHS produces a noticeable improvement not only in the long-term performance of recurrent truncation selection, but also will surpass the genetic gain of a population that did not have that single generation of OHS within 10 generations. The hybrid selection approach outperforms high-genetic-gain OCS in the first 40 generations of selection, though it does exhaust population diversity faster than OCS.

This study’s results advise that OHS could be investigated further for practical scenarios characterised by extracting useful selections from diverse candidate pools with large breeding value differences, or recurrent selection on small candidate pools. OHS could be suited to early generations of an elite breeding program after the introduction of new diversity (to take best advantage of the new diversity and beneficial recombinants), in resistance breeding (for stacking multiple resistances together), or in pre-breeding programs that explore genetic resources from gene banks (as it can take into account useful diversity at block level), though further investigation would be required to find how to best apply the method in these practical scenarios.

This study therefore shows that this evolutionary computing-based haplotype stacking approach to selection is a promising avenue for further investigation.

## Acknowledgements

The authors are grateful for funding from the Australian Research Council, Linkage Project LP170100317, “FastStack - evolutionary computing to stack desirable alleles in wheat”.

## Data Availability

The raw or processed simulated datasets generated during the current study are available from the corresponding author on request.

## Abbreviations

GEBV: genomic estimated breeding value
GS: genomic selection
OCS: optimal cross selection
OHS: optimal haplotype selection
OPV: optimal population value
SNP: single nucleotide polymorphism

